# Encoding Cell Phenotype from Label-Free Imaging Flow Cytometry with Unsupervised Deep Learning

**DOI:** 10.1101/2025.05.29.656720

**Authors:** Benjamin Gincley, Farhan Khan, Ananya Kumar, Miguel Fuentes-Cabrera, Jeremy Guest, Ameet J. Pinto

## Abstract

Phenotype characterization with single-cell resolution can enable deep and nuanced insights into microbiological systems. Currently, Flow Cytometry and Imaging Flow Cytometry (IFC) offer numerous advantages, but are marred by barriers to accessibility: (1) high instrument costs; (2) labor-intensive, technically demanding sample preparation; and (3) reliance on consumable reagents (i.e., fluorescent labels). To achieve phenotype characterization without these constraints, we evaluated the low-cost, low-input ARTiMiS IFC as a potential alternative instrument technology. To demonstrate this approach, we used intracellular lipid content in microalgae, an important phenotype for production of biofuels and high-value bioproducts, as the phenotype of interest. Variational Auto-Encoder (VAE) unsupervised deep learning methodology was implemented to encode phenotype variation from un-annotated training data. The VAE embeddings were compared with other label-free predictor modalities to evaluate the stability of VAE data encoding across replicates and its predictive power to estimate the target phenotype. The VAE embeddings were robust and consistent between culture batches, and yielded accurate, consistent predictions of the demonstration phenotype in a high-throughput, non-destructive, dye-free methodology. In this proof-of-concept study, we demonstrate that VAE-enabled ARTiMiS IFC may serve as a viable alternative for cell phenotype characterization while overcoming several of the key drawbacks of traditional high-fidelity techniques.

**Synopsis:** Label-free Imaging Flow Cytometry data was processed by a Variational Auto-Encoder to accurately predict lipid content in microalgal cells.

## Introduction

Flow Cytometry (FCM) has been used extensively for cell phenotyping in wide range of applications ranging from immunophenotyping,^1^ cell viability assessment^2^ to cellular function characterization^3^. The popularity of FCM has been driven by numerous hardware and software improvements for over 50 years, as well as standardization of fluorescent labels, protocols, and analysis workflows in disparate fields from immunology to environmental microbiology that have collectively contributed to a mature, reliable technology.^4,5^ However, FCM has several limitations that hinder its widespread adoption such as high instrumentation costs,^6^ reliance on fluorescent labeling,^7^ and specialized training requirements.^8^ Imaging Flow Cytometry (IFC), also known as Flow Imaging Microscopy (FIM), is another versatile, high-throughput technology that can provide single-cell resolution.^9^ As with FCM, fluorescent staining has dramatically improved quantitative cell analysis using IFC,^10^ though reliance on cell labeling imposes its own set of restrictions. Namely, large-scale labeling assays are labor-intensive and expensive, and incomplete knowledge of the markers associated with undiscovered or poorly-understood cell populations can preclude label selection and thus characterization.^11^ These obstacles have motivated the exploration of label-free IFC technologies and techniques to overcome such challenges.

Microscopic image analysis has benefited from advancements in computer vision and deep learning, fostering new insights into cell phenotyping and physiology.^12,13^ Recent developments in deep learning may provide new techniques to analyze microscope image data, particularly to enable quantitative label-free phenotyping using IFC. While deep convolutional neural network (CNN) models have been widely reported for cell classification tasks, such models require, at a minimum, “weak” supervision for training.^14^ In applications where physiological conditions or cell states are not discrete or well understood, implementation of accurate supervised models is not possible. In contrast, variational auto-encoders (VAEs)^15,16^ that encode visual information into a low-dimensional coordinate space (referred to as “latent space”) demonstrate potential to infer continuous phenotypes from uncategorized training data.^17,18^ Importantly, phenotypic variation can be encoded without either (1) *a priori* awareness of the visual signatures corresponding to specific cell states of interest (“strong” supervision), or (2) the condition state of individual samples comprising the training data pool (“weak” supervision). This unsupervised approach may therefore dramatically improve accessibility and automation by reducing requirements for exhaustive sample knowledge and well-annotated observations to curate data for model training.

Recent studies^18–20^ have developed workflows for cell phenotype characterization using VAE models, but almost exclusively rely on sophisticated imaging technologies (e.g., laser-scanning confocal microscopy) which does little to address the previously cited obstacles to broader adoption of FCM. To date, there has not been a systematic investigation into whether low-cost imaging platforms developed for environmental applications such as PlanktoScope^21^ or ARTiMiS^22^ can capture images of sufficient detail to train robust and accurate VAEs. In this study, we assess whether robust VAEs can be trained from label-free ARTiMiS image data to encode an industrially relevant cell phenotype. As a proof-of-concept, we investigated an important phenotype in microalgal biomass applications: intracellular lipid content. High lipid microalgal biomass is an important feedstock for conversion into biofuels (e.g., sustainable aviation fuels) for commodity markets, or high-value bioproducts bound for consumer markets including omega fatty acids such as DHA.^23,24^ Additionally, the explicit cultivation conditions that drive lipid accumulation have been well-documented,^25,26^ making this an ideal model system to test this approach. In this work, we explored the effectiveness of VAE-enabled ARTiMiS IFC to predict an environmentally relevant phenotype without labeling dyes, extensive sample preparation protocols, or capital-intensive instrumentation, utilizing dye-labeled flow cytometry data as a ground-truth reference.

## Materials and Methods

### Cell Culture and Conditions

*Chlorella sorokiniana* (UTEX 1602) was obtained from the University of Texas at Austin algae culture collection and maintained in Bold’s 1NV medium at 25°C under a 12h:12h day-night cycle with a photosynthetically active radiation (PAR) value of 80 µmol photons m^-2^·s^-1^ and 100 rpm agitation. Nitrogen (N) limitation was enforced to induce variation in lipid content phenotype.^26–28^ *C. sorokiniana* was maintained in two distinct conditions: N-replete standard Bold’s growth medium (i.e., control), and N-deplete modified Bold’s medium lacking nitrate to drive greater lipid accumulation relative to control. Paired batch culture flasks were seeded from the same N-replete maintenance culture, split into N-replete (NR) (control) and N-deplete (ND) (induced phenotype) media for cultivation during the experimental period.

This study was performed in two phases. Phase 1: batches were cultivated as synchronous biological duplicates (referred to as Batches 1A and 1B) in 125 mL Erlenmeyer shake flasks with 40 mL starting volume and maintained for 7 days. Phase 2: a third, asynchronous biological replicate (referred to as Batch 2) batch culture was maintained in 500 mL Erlenmeyer shake flasks with 150 mL starting volume and cultivated for 21 days. The Phase 2 experiment was conducted to determine whether results remained consistent in subsequent generations of the same cell line.

### Flow Cytometry

Samples were stained with the lipophilic fluorescent dye BODIPY 505/515^27,29,30^ and processed on a CytoFLEX flow cytometer (Becton Dickinson Life Sciences, USA) to determine intracellular lipid content. Events were identified as viable microalgae by size characteristics (forward scatter, FSC; side scatter, SSC), autofluorescence (Blue excitation laser, 690/50 nm bandpass emission; Red laser, 660/10 nm emission), and lipophilic dye signal (Blue laser, 525/40 nm emission; later referred to by channel alias B525). Detected event emission area (e.g., B525-A) measurements were used for all channels. Samples were prepared as technical triplicates, processed to a stopping criterion of 15,000 events, and aggregated during post-processing analysis. While the overlap between microalgal pigment autofluorescence and BODIPY 505/515 emission has been ignored previously,^27,30,31^ physiological state can cause shifts in autofluorescence spectra that may interfere with BODIPY 505/515 emission measurements.^32^ In this study, microalgae pigment autofluorescence spillover into the 525/40 nm channel was low yet measurable. Thus, compensation was calculated from linear regression of unlabeled cells^33^ to negate the impact of pigment content on lipid measurements. Compensation regressions were fit per sample run (day) from unlabeled cells from each condition (ND and NR) and subtracted from their respective raw fluorescence measurements by applying the compensation (Equation 1) on a per-event (*i*) basis, where *a* and *b* represent coefficients of the line fitting for the respective unlabeled control.

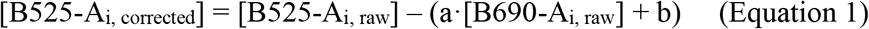

### ARTiMiS Imaging Flow Cytometry

Samples were processed in parallel on FCM and on the ARTiMiS IFC system using the previously described standard instrument configuration.^34^ Briefly, a samples were diluted with respective medium to cell concentrations of 10^5^-10^6^ cells·mL^-1^ and a settling time of 90 seconds was chosen to maximize the proportion of cells in the same focal plane for each frame. “Low-quality” regions of interest (ROIs) – multi-object ROIs, non-cell objects (bubbles, debris), and out-of-focus objects – were screened using empirically determined feature filters and removed. For a complete list of filter criteria, see **Table S1**. ROIs were subsequently manually reviewed for quality control. All those passing quality control were included for model training or inference.

### Variational Auto-Encoder Deep Learning Model

A VAE model was trained to encode the complete spectrum of phenotypic variation exhibited during the observation period. The complete Batch 1A image dataset (all conditions and time points of N-Deplete 1A and N-Replete 1A) was train/test split (80% for model training, 20% holdout for evaluation). The training set was augmented 8-fold through a complete set of rotations and mirrors to increase dataset size (previously described)^22,34^ for a total of 119,136 training images (13,992 unique images). The VAE model architecture was adapted from the “micro-CNN” design previously reported^22,34^ to be well-suited for ARTiMiS-generated image data (TensorFlow v2.5.0, Python). For a complete description of the VAE architecture used, see **Table S2**. VAEs were trained on an NVIDIA RTX 3070 Ti (8GB VRAM) GPU for 12 epochs (until loss did not further decrease). VAEs were iteratively trained and screened for optimal fit to test regression data. A diagnostic score was calculated from the root mean squared error (RMSE) of linear regression fitting of the Batch 1A test data embedding to Batch 1A flow cytometry measurements, effectively scoring how well a particular VAE’s embedding represented the complementary flow cytometry data. After identifying a leading VAE model, the same model, without re-training, was then used to perform inference on unseen sample data from Batch 1B and Batch 2, yielding latent space coordinate embeddings for each ROI. These embeddings were used for all VAE-related analysis.

### Statistical Evaluation of VAE Embedding Coordinates

To evaluate population statistics of VAE model embeddings, Spearman’s rank correlation coefficient was used to determine correlation between embedding coordinates and sampling day, as well as ARTiMiS-measured morphological features. Euclidean distance used to evaluate distance between populations. One-way ANOVA was used to evaluate statistical significance of differences in population mean values.

### Phenotype Prediction Models and Evaluation

Despite FCM and IFC both having single-cell resolution, it was not possible for measurements from both instruments to be paired at a single-cell level; rather, only sample population means could be used for comparison. Constrained by the number of samples (n=36) to perform regression, linear regression and random forest regression were selected. Predictor features (i.e., 25 morphological feature measurements directly from ARTiMiS, two latent dimensions from VAE, and two physical FCM measurements – FSC, SSC) were dimensionally reduced using principal component analysis (PCA) to compare the impact of number of features on prediction performance. During evaluation, paired data were 80:20 train/test split with random reshuffling (Monte Carlo cross validation) for 500 iterations prior to model fitting to reduce outlier effects. Mean values with standard deviation from fitting iterations were reported for coefficient of determination R^2^ and root mean squared error (RMSE).

## Results and Discussion

### Nitrogen limitation resulted in lipid accumulation in C. sorokiniana

In agreement with prior studies,^26–28,35^ flow cytometric analysis of intracellular lipid content labeled with the fluorescent lipophilic dye BODIPY 505/515 indicated negligible lipid accumulation in the NR condition during the study period. In contrast, the ND condition exhibited a significant accumulation of intracellular lipids (**Figure 1A-B**). In all three batches, lipid levels peaked five days after inoculation, then decreased slightly by Day 7. During the longer observation period in Batch 2 (**Figure 1B**), lipid content increased after Day 7 in both NR and ND conditions with increase in lipid content in NR condition attributable to nitrogen limitation after day 7. While mean fluorescence intensity (MFI) results (**Figure 1A-B**) imply a stark difference between cell populations of the two conditions, population distributions of individual sampling days, for example Day 7 (**Figure 1C-D**), reveal heterogeneous populations exhibiting a wide range of lipid content phenotypes.

**Figure 1.**
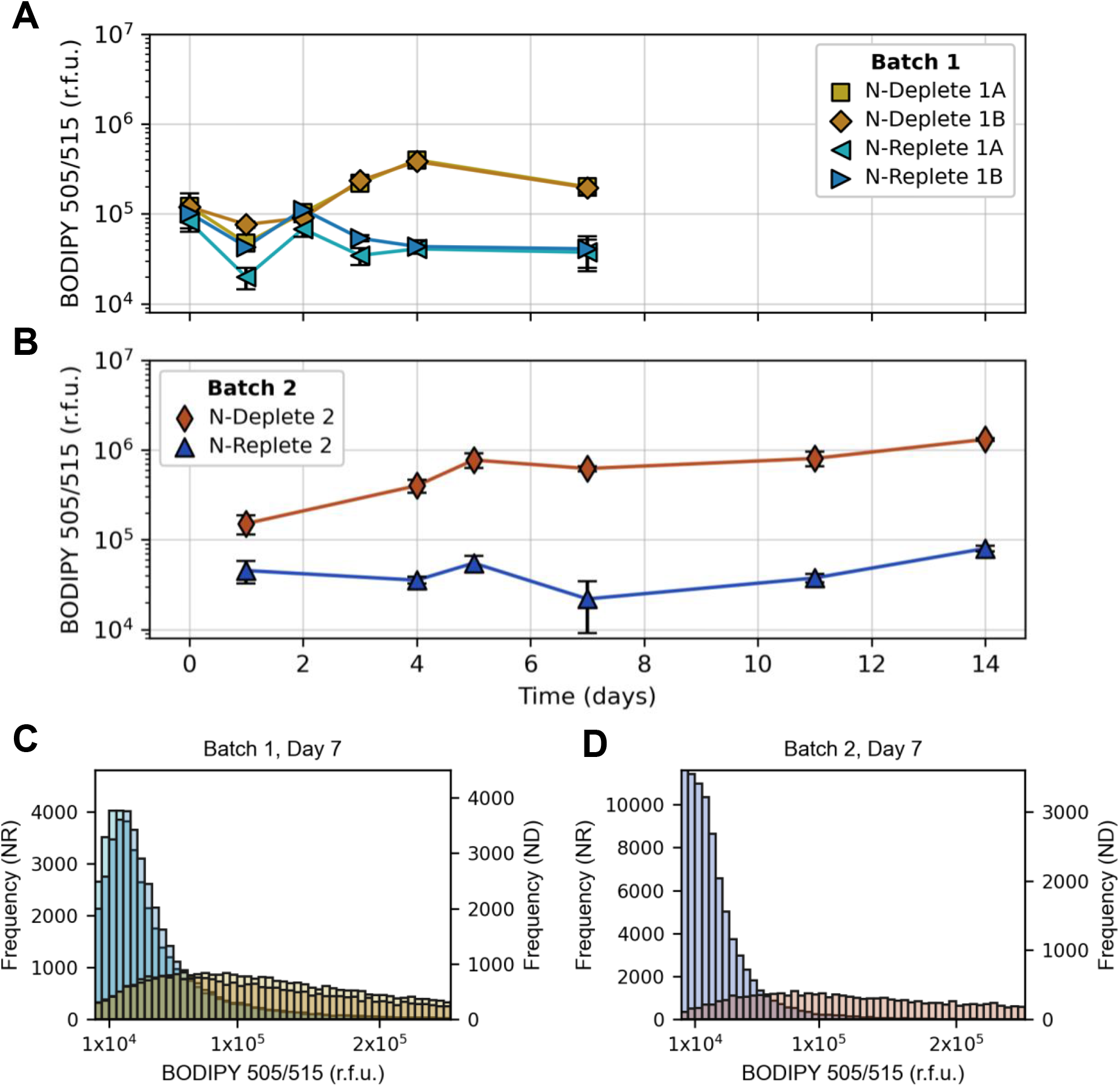
Mean Fluorescence Intensity (MFI, relative fluorescence units) of lipophilic dye BODIPY 505/515 during batch cultivation in ND and NR media conditions. **(A)** Samples in Batches 1A and 1B were cultured in parallel, **(B)** Batch 2 included independent replication of ND and NR conditions for a longer observation period. Error bars indicate standard deviation of population means from **(A)** 3 and **(B)** 4 technical replicates per condition, respectively. **(C**,**D)** Event histograms from Day 7 samples depict the heterogeneity of BODIPY signal, particularly among ND populations. ND and NR conditions are colored as in A, B.

### Variational Auto-Encoder training yielded reproducible embeddings

VAEs efficiently reduce the dimensional complexity of image data (e.g., a 70×70 pixel ROI having 4900 “dimensions”) to a significantly more manageable 1-3 dimensions in “latent” space.^16^ After training a VAE on the image data of one batch (i.e., Batch 1A), the synchronous (i.e., Batch 1B) and asynchronous (i.e., Batch 2) replicate pairs were used to evaluate the “stability” of the embedding. An embedding would be considered “stable” if evaluation samples were mapped to the same coordinate regions as the training samples for a given phenotype state. Embedding stability is an important attribute for such a VAE model as it would mean that training could be performed once from a diagnostic dataset, then the same model could be used on future batches without retraining.

A VAE model was trained using 80% of the image data collected from Batch 1A. Based on the flow cytometry reference results, these samples spanned a range of phenotype states from low to high intracellular lipid content. The remaining 20% holdout data fraction was used to probe the latent space after training (**Figure 2A**). By labeling individual points with their nutrient condition and time point, it was possible to examine the latent space to detect patterns that would imply an orderly (or disorderly) structure to the coordinate system. The first latent dimension (LD1) appeared to encode temporal information: sample date was seen to increase with increasing LD1 coordinate (magenta arrow). Populations were grouped by sample condition and evaluated for statistical significance via Spearman correlation. All conditions independently exhibited a significant (p < 0.05) correlation between sample day and LD1, with correlation coefficients ranging from 0.61 (ND 1A and 1B) to 0.09 (NR 2). To further evaluate the statistical stability of the embedding, sample conditions were grouped by day to determine the consistency of mean latent coordinates via ANOVA. Of the 22 sample pairs synchronized by day (e.g., ND 1A Day 1 and ND 1B Day 1; NR 1B Day 7 and NR 2 Day 7), there were 9 instances of indistinguishable mean coordinates for at least one dimension (LD1 or LD2). Among those, one pair (Day 7 NR, 1A and 1B) shared the same mean coordinate on both dimensions, and Day 7 ND samples shared the same LD1 coordinate for all three replicates. These results indicated that the VAE model was stable across different culture batches and that a model could be trained once and remain accurate for sequential batches without retraining or recalibration.

**Figure 2.**
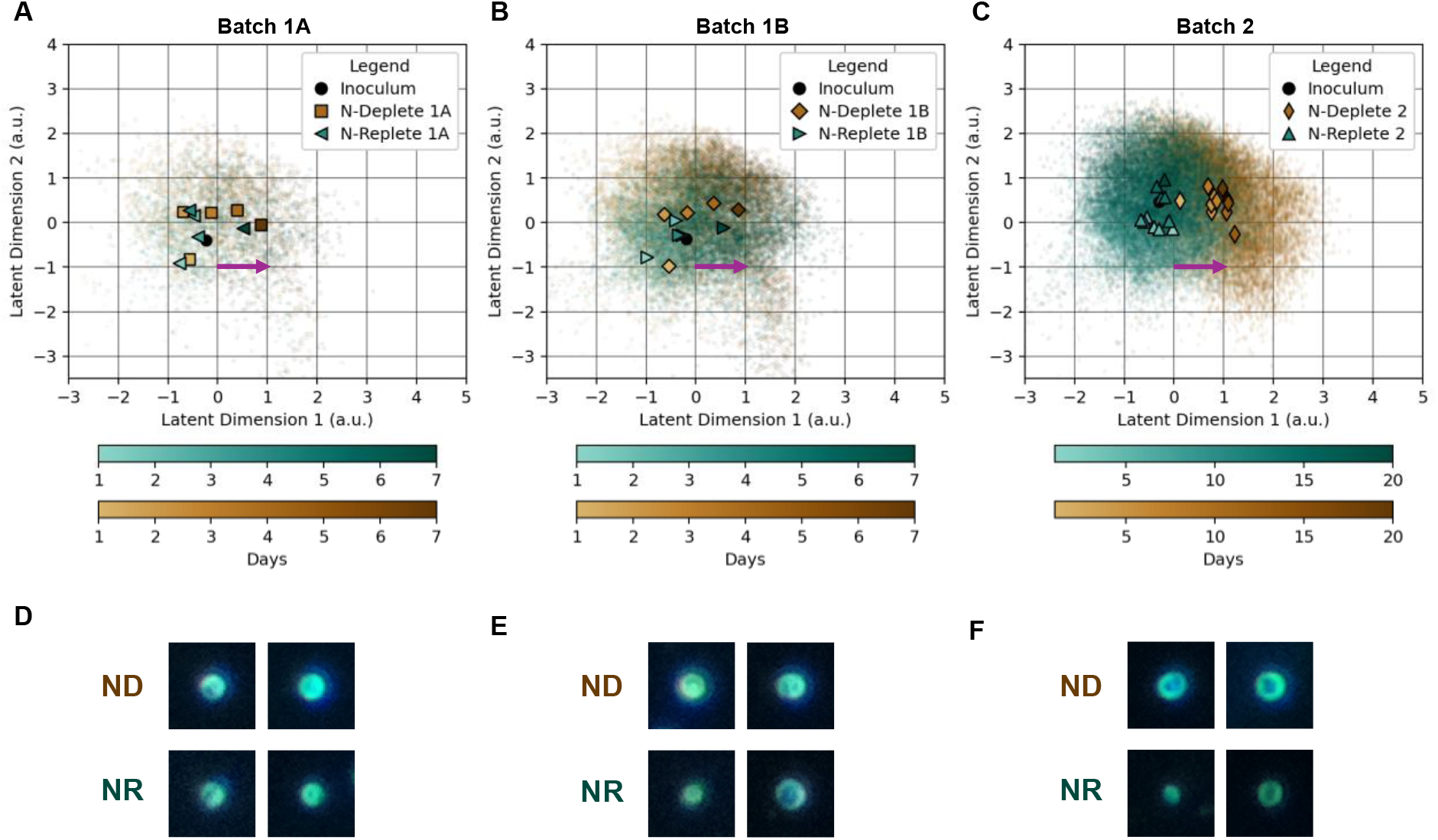
Latent space encodings of **(A)** the test set from Batch 1A (20% of total events) and the evaluation batches **(B)** Batch 1B and **(C)** Batch 2 (100% of events). Scatter points depict individual events, prominent markers indicate mean coordinates for each sample time point (days) for each condition. Color value scales by observation day, darker values corresponding to later days. Note the reduced population size of **(A)** is due to the test holdout comprising 20% of the total dataset. For Batch 1, samples were collected on Days 0, 1, 2, 3, 4, 7. For Batch 2, samples were collected on Days 0, 1, 4, 5, 6, 7, 8, 11, 13, 14, 15, 20. Magenta arrows indicate coordinate procession as lipid content increases. **(D, E, F)** Example darkfield images of events (cells) observed on Day 7 of Batches 1A, 1B, and 2, respectively, from conditions N-Deplete (above) and N-Replete (below).

Batch 2 exhibited a greater separation between conditions than Batch 1 samples (**Figure 2C**). The global centroid of all points for each condition was measured for comparison. The distances between the centroids of ND 1A and NR 1A samples, and ND 1B and NR 1B samples, were 0.30 and 0.39, respectively, whereas the distance between ND 2 and NR 2 centroids was 1.16. The same pattern was observed when analyzing lipid content data from FCM (**Figure 1**) – Batch 1 mean fluorescence intensity (MFI) values were closer in magnitude, particularly for Days 0-2, whereas Batch 2 MFI measurements were significantly distant as early as Day 1.

### VAE primarily encoded features relating to cell size and edge sharpness

To gain a more intuitive understanding of the phenotypic meaning of this coordinate space, an investigation into which visual features were being encoded was performed. After training, the VAE could be qualitatively interpreted by probing its decoder component, which up-samples a given embedding coordinate back into a full-size image. This process generated archetypal images spanning the coordinate system (**Figure S1**), providing an intuitive visualization of the types of morphological variations corresponding to specific coordinate regions. A correlation analysis was performed between embedding coordinates and morphological feature data derived from traditional computer vision image processing (**Figure S2**) for a quantitative characterization. Overall, features relating to cell size and edge sharpness changed as a function of embedding coordinate, congruent with the visualized decoder output (**Figure S1**). This finding aligns with recent studies observing morphological changes in nitrogen-stressed microalgae.^36,37^

### VAE embeddings translated to robust phenotype estimates

While there was an established pattern underpinning the embedding coordinates, the effectiveness of using such embeddings to predict a target cell phenotype required evaluation to determine if use of VAEs provided a measurable improvement over alternative methods. To that end, the efficiency of translating VAE model outputs into a predictive model for cell phenotype was tested against two alternatives: (1) the use of traditional image data processing outputs – geometric morphological features from standard computer vision techniques, and (2) the use of non-fluorescent measurements from flow cytometry, specifically the forward scatter (FSC) and side scatter (SSC) channels. For this assessment, multiple linear regression, simple linear regression (after applying dimensionality reduction via principal component analysis), and random forest regression were used to fit the proxy method measurements to BODIPY 505/515 FCM measurements. Model fit coefficient of determination R^2^ and root mean squared error (RMSE) were selected to compare performance. Regression models were iteratively fit from 500 cross validation iterations, randomly reshuffling 80% training, 20% testing data splits to provide representative average performance results from 36 unique samples.

Random forest regression from FCM physical measurements (FSC and SSC) produced the best overall approximation for BODIPY 505/515 measurements in absolute terms (significantly lower RMSE than all other models, **Table S3**). However, the PCA-reduced 1D simple linear regression from the VAE embedding was the next best performing (lowest RMSE) model, followed closely by the 2D feature table data random forest regression and 2D VAE linear regression (**Figure 3, Table S3**). These findings (along with the coupled R^2^ values from these models) indicated that given the correct dimensionality reduction and regression model selection, each modality: FCM, VAE, and feature table data, could be used to approximate the target phenotype with near-equivalent accuracy. However, among the permutations tested, the VAE-based approximations most consistently outperformed the other models in the set (ranking 2^nd^, 3^rd^, and 5^th^ of the top 5 methods for lowest RMSE, 3^rd^ and 4^th^ for R^2^). Thus, VAE embedding data as predictor features were the least sensitive to pre-processing methods and regression model type. In comparison, feature table data and physical FCM channel measurements as predictor variables often yielded significantly worse approximations in the “wrong” configurations.

**Figure 3.**
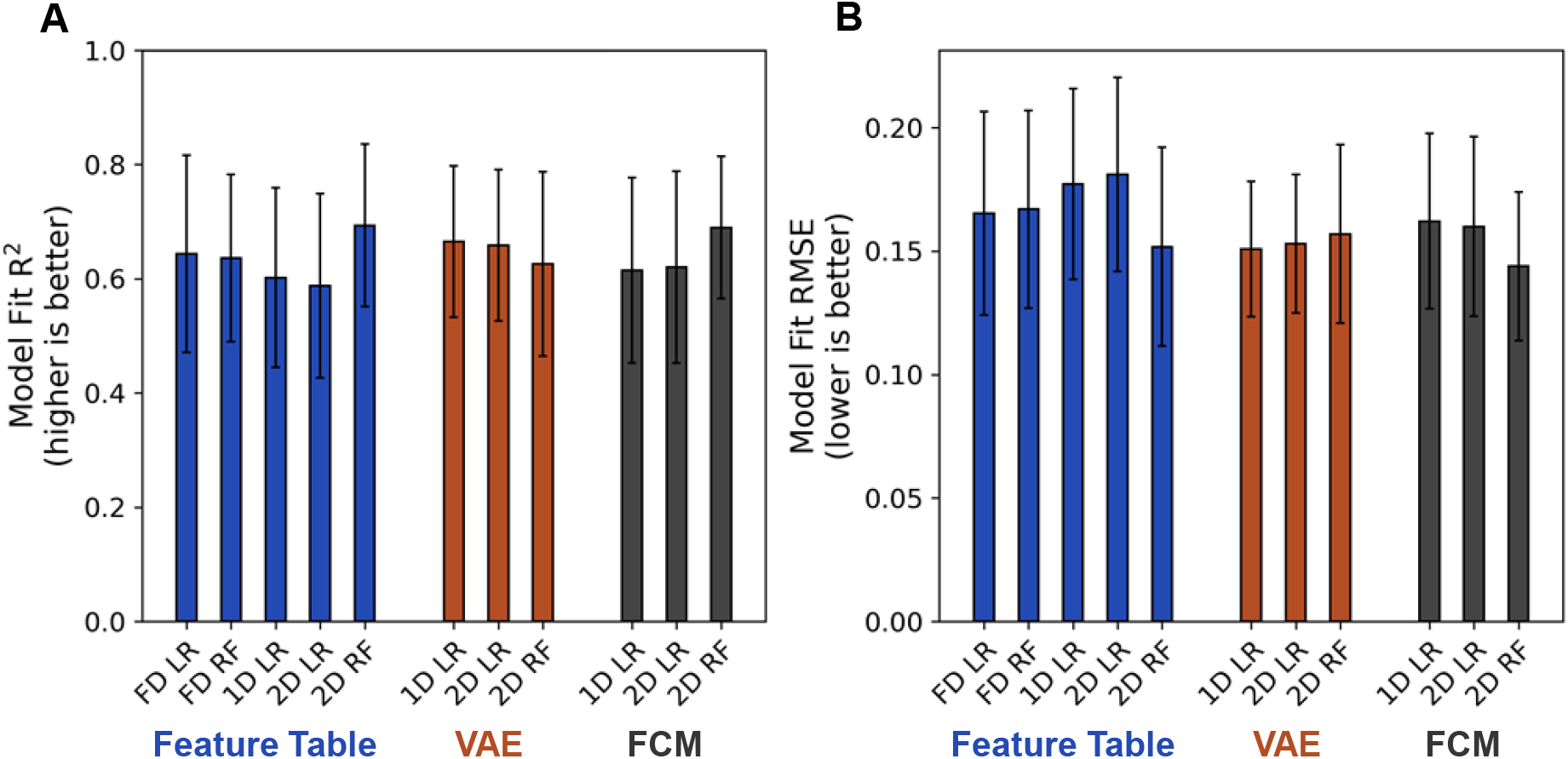
Regression scores of (A) coefficient of determination, R^2^, and (B) root mean squared error, RMSE, evaluated from fitting: feature table data directly from ARTiMiS (blue), VAE embedding coordinate data (orange) and label-free flow cytometry measurements (black) to predict BODIPY 505/515 measurements. FD indicates fully-dimensioned feature table data (25 features), 1D and 2D indicate 1-dimension and 2-dimension predictor data, respectively. For dimensionalities less than native for the type (e.g., 2D for Feature Table, 1D for VAE, but not 2D for VAE or FCM, etc.), dimensionality was reduced via principle component analysis. Values and error bars report mean and standard deviation, respectively, from 500 cross-validation iterations of 36 total sample points split 80:20 for fitting and evaluation. Pairwise significance levels for RMSE depicted in Table S3.

These results provided strong evidence that label-free FCM (using only FSC and SSC channels) was effective at approximating this phenotype, applicable in situations where fluorescent labeling is not possible. However, such situations where FCM is available but fluorescent labeling is not, are expected to be uncommon. A more probable context is the need to provide label-free phenotype estimates without access to capitally intensive instrumentation, perishable reagents, and skilled labor. These findings suggest that IFC can be effective at collecting relevant data for phenotype estimation, and that transforming raw image data into latent embeddings with VAE models can provide more consistent and robust predictor features compared to traditional computer vision image processing techniques.

These data were collected on the low-cost ARTiMiS IFC system. While this is not the first study to report the use of VAEs for cell phenotype characterization, previous studies have relied on sophisticated instruments, such as confocal microscopes, for data collection where samples were fluorescently labeled.^17–19^ In the few instances of label-free cell characterization with VAEs, techniques such as quantitative phase imaging (QPI)^28^ or phase contrast imaging in a controlled environment chamber^38^ have been employed. Uniquely, this is the first study to demonstrate application of VAEs for label-free imaging on a low-cost, portable imaging flow cytometry platform. While the lipid content in microalgae was selected for method demonstration, this approach has the potential to work in other applications where small variations in cell morphology may indicate changes in a target phenotype. VAE-enabled label-free IFC represents a promising new technique to reduce the capital and technical burdens of cell characterization. Widespread adoption of this approach could ultimately translate to lower operational costs and potentially greater yields for industrial biological processes.

## Acknowledgement

This work was funded by the U.S. Department of Energy, Office of Energy Efficiency and Renewable Energy, under Award Number DE-EE0009270. This work was also supported by the U.S. National Science Foundation Graduate Research Fellowship Program under Award Number 2039655 and the Paul L Busch Award for Innovation in Applied Water Quality Research. Any opinions, findings, and conclusions or recommendations expressed in this publication are those of the author(s) and do not necessarily reflect the views of the U.S. Department of Energy.

## References

(1) Pockley, A. G.; Foulds, G. A.; Oughton, J. A.; Kerkvliet, N. I.; Multhoff, G. Immune Cell Phenotyping Using Flow Cytometry. Current Protocols in Toxicology 2015, 66 (1), 18.8.1-18.8.34. 10.1002/0471140856.tx1808s66.

(2) Weaver, J. L. Estimation of Cell Viability by Flow Cytometry. In Flow Cytometry Protocols; Jaroszeski, M. J., Heller, R., Eds.; Humana Press: Totowa, NJ, 1998; pp 77–83. 10.1385/0-89603-354-6:77.

(3) Bleesing, J. J. H.; Fleisher, T. A. Cell Function-Based Flow Cytometry. Seminars in Hematology 2001, 38 (2), 169–178. 10.1016/S0037-1963(01)90050-2.

(4) Robinson, J. P.; Ostafe, R.; Iyengar, S. N.; Rajwa, B.; Fischer, R. Flow Cytometry: The Next Revolution. Cells 2023, 12 (14), 1875. 10.3390/cells12141875.

(5) Czechowska, K.; Johnson, D. R.; van der Meer, J. R. Use of Flow Cytometric Methods for Single-Cell Analysis in Environmental Microbiology. Current Opinion in Microbiology 2008, 11 (3), 205–212. 10.1016/j.mib.2008.04.006.

(6) Brestoff, J. R.; Frater, J. L. Contemporary Challenges in Clinical Flow Cytometry: Small Samples, Big Data, Little Time. The Journal of Applied Laboratory Medicine 2022, 7 (4), 931–944. 10.1093/jalm/jfab176.

(7) MacLennan, C. A.; Liu, M. K. P.; White, S. A.; Oosterhout, J. J. G. van; Simukonda, F.; Bwanali, J.; Moore, M. J.; Zijlstra, E. E.; Drayson, M. T.; Molyneux, M. E. Diagnostic Accuracy and Clinical Utility of a Simplified Low Cost Method of Counting CD4 Cells with Flow Cytometry in Malawi: Diagnostic Accuracy Study. BMJ 2007, 335 (7612), 190–190. 10.1136/bmj.39268.719780.BE.

(8) Muccio, V. E.; Saraci, E.; Gilestro, M.; Oddolo, D.; Ruggeri, M.; Caltagirone, S.; Bruno, B.; Boccadoro, M.; Omedè, P. Relevance of Sample Preparation for Flow Cytometry. International Journal of Laboratory Hematology 2018, 40 (2), 152–158. 10.1111/ijlh.12755.

(9) Rees, P.; Summers, H. D.; Filby, A.; Carpenter, A. E.; Doan, M. Imaging Flow Cytometry. Nat Rev Methods Primers 2022, 2 (1), 1–13. 10.1038/s43586-022-00167-x.

(10) Tan, W. C. C.; Nerurkar, S. N.; Cai, H. Y.; Ng, H. H. M.; Wu, D.; Wee, Y. T. F.; Lim, J. C. T.; Yeong, J.; Lim, T. K. H. Overview of Multiplex Immunohistochemistry/Immunofluorescence Techniques in the Era of Cancer Immunotherapy. Cancer Communications 2020, 40 (4), 135–153. 10.1002/cac2.12023.

(11) Lee, K. C. M.; Wang, M.; Cheah, K. S. E.; Chan, G. C. F.; So, H. K. H.; Wong, K. K. Y.; Tsia, K. K. Quantitative Phase Imaging Flow Cytometry for Ultra-Large-Scale Single-Cell Biophysical Phenotyping. Cytometry Part A 2019, 95 (5), 510–520. 10.1002/cyto.a.23765.

(12) Pratapa, A.; Doron, M.; Caicedo, J. C. Image-Based Cell Phenotyping with Deep Learning. Current Opinion in Chemical Biology 2021, 65, 9–17. 10.1016/j.cbpa.2021.04.001.

(13) Doan, M.; Barnes, C.; McQuin, C.; Caicedo, J. C.; Goodman, A.; Carpenter, A. E.; Rees, P. Deepometry, a Framework for Applying Supervised and Weakly Supervised Deep Learning to Imaging Cytometry. Nat Protoc 2021, 16 (7), 3572–3595. 10.1038/s41596-021-00549-7.

(14) Caicedo, J. C.; McQuin, C.; Goodman, A.; Singh, S.; Carpenter, A. E. Weakly Supervised Learning of Single-Cell Feature Embeddings; Proceedings of the IEEE Conference on Computer Vision and Pattern Recognition (CVPR), 2018; pp 9309–9318.

(15) Kingma, D. P.; Welling, M. Auto-Encoding Variational Bayes. In 2nd International Conference on Learning Representations, ICLR 2014, Banff, AB, Canada, April 14-16, 2014, Conference Track Proceedings; Bengio, Y., LeCun, Y., Eds.; 2014.

(16) Kingma, D. P.; Welling, M. An Introduction to Variational Autoencoders. arXiv.org. 10.1561/2200000056.

(17) Ternes, L.; Dane, M.; Gross, S.; Labrie, M.; Mills, G.; Gray, J.; Heiser, L.; Chang, Y. H. A Multi-Encoder Variational Autoencoder Controls Multiple Transformational Features in Single-Cell Image Analysis. Commun Biol 2022, 5 (1), 1–10. 10.1038/s42003-022-03218-x.

(18) Chow, Y. L.; Singh, S.; Carpenter, A. E.; Way, G. P. Predicting Drug Polypharmacology from Cell Morphology Readouts Using Variational Autoencoder Latent Space Arithmetic. PLOS Computational Biology 2022, 18 (2), e1009888. 10.1371/journal.pcbi.1009888.

(19) Burgess, J.; Nirschl, J. J.; Zanellati, M.-C.; Lozano, A.; Cohen, S.; Yeung-Levy, S. Orientation-Invariant Autoencoders Learn Robust Representations for Shape Profiling of Cells and Organelles. Nat Commun 2024, 15 (1), 1022. 10.1038/s41467-024-45362-4.

(20) Sandarenu, P.; Chen, J.; Slapetova, I.; Browne, L.; Graham, P. H.; Swarbrick, A.; Millar, E. K. A.; Song, Y.; Meijering, E. Semi-Supervised Variational Autoencoder for Cell Feature Extraction In Multiplexed Immunofluorescence Images. In 2024 IEEE International Symposium on Biomedical Imaging (ISBI); 2024; pp 1–5. 10.1109/ISBI56570.2024.10635107.

(21) Pollina, T.; Larson, A. G.; Lombard, F.; Li, H.; Le Guen, D.; Colin, S.; de Vargas, C.; Prakash, M. PlanktoScope: Affordable Modular Quantitative Imaging Platform for Citizen Oceanography. Frontiers in Marine Science 2022, 9. 10.3389/fmars.2022.949428.

(22) Gincley, B.; Khan, F.; Alam, M. M.; Hartnett, E.; Kim, G.-Y.; Molitor, H. R.; Fisher, A.; Bradley, I.; Guest, J.; Pinto, A. J. Morphotype-Resolved Characterization of Microalgal Communities in a Nutrient Recovery Process with ARTiMiS Flow Imaging Microscopy. Water Research 2025, 123801. 10.1016/j.watres.2025.123801.

(23) Barkia, I.; Saari, N.; Manning, S. R. Microalgae for High-Value Products Towards Human Health and Nutrition. Mar Drugs 2019, 17 (5), 304. 10.3390/md17050304.

(24) Qu, L.; Ren, L.-J.; Huang, H. Scale-up of Docosahexaenoic Acid Production in Fed-Batch Fermentation by Schizochytrium Sp. Based on Volumetric Oxygen-Transfer Coefficient. Biochemical Engineering Journal 2013, 77, 82–87. 10.1016/j.bej.2013.05.011.

(25) An, M.; Gao, L.; Zhao, W.; Chen, W.; Li, M. Effects of Nitrogen Forms and Supply Mode on Lipid Production of Microalga Scenedesmus Obliquus. Energies 2020, 13 (3), 697. 10.3390/en13030697.

(26) Bono, M.S. (Jr.); Garcia, R. D.; Sri-Jayantha, D. V.; Ahner, B. A.; Kirby, B. J. Measurement of Lipid Accumulation in Chlorella Vulgaris via Flow Cytometry and Liquid-State 1H NMR Spectroscopy for Development of an NMR-Traceable Flow Cytometry Protocol. PLOS ONE 2015, 10 (8), e0134846. 10.1371/journal.pone.0134846.

(27) Benito, V.; Goñi-de-Cerio, F.; Brettes, P. BODIPY Vital Staining as a Tool for Flow Cytometric Monitoring of Intracellular Lipid Accumulation in Nannochloropsis Gaditana. J Appl Phycol 2015, 27 (1), 233–241. 10.1007/s10811-014-0310-x.

(28) Chen, C. L.; Mahjoubfar, A.; Tai, L.-C.; Blaby, I. K.; Huang, A.; Niazi, K. R.; Jalali, B. Deep Learning in Label-Free Cell Classification. Sci Rep 2016, 6 (1), 21471. 10.1038/srep21471.

(29) Koreivienė, J. Microalgae Lipid Staining with Fluorescent BODIPY Dye. In Biofuels from Algae: Methods and Protocols; Spilling, K., Ed.; Methods in Molecular Biology; Springer: New York, NY, 2020; pp 47–53. 10.1007/7651_2017_101.

(30) Rumin, J.; Bonnefond, H.; Saint-Jean, B.; Rouxel, C.; Sciandra, A.; Bernard, O.; Cadoret, J.-P.; Bougaran, G. The Use of Fluorescent Nile Red and BODIPY for Lipid Measurement in Microalgae. Biotechnology for Biofuels 2015, 8 (1), 42. 10.1186/s13068-015-0220-4.

(31) Südfeld, C.; Hubáček, M.; D’Adamo, S.; Wijffels, R. H.; Barbosa, M. J. Optimization of High-Throughput Lipid Screening of the Microalga Nannochloropsis Oceanica Using BODIPY 505/515. Algal Research 2021, 53, 102138. 10.1016/j.algal.2020.102138.

(32) Cheloni, G.; Slaveykova, V. I. Optimization of the C11-BODIPY581/591 Dye for the Determination of Lipid Oxidation in Chlamydomonas Reinhardtii by Flow Cytometry. Cytometry Part A 2013, 83 (10), 952–961. 10.1002/cyto.a.22338.

(33) Roca, C. P.; Burton, O. T.; Gergelits, V.; Prezzemolo, T.; Whyte, C. E.; Halpert, R.; Kreft, Ł.; Collier, J.; Botzki, A.; Spidlen, J.; Humblet-Baron, S.; Liston, A. AutoSpill Is a Principled Framework That Simplifies the Analysis of Multichromatic Flow Cytometry Data. Nat Commun 2021, 12 (1), 2890. 10.1038/s41467-021-23126-8.

(34) Gincley, B.; Khan, F.; Hartnett, E.; Fisher, A.; Pinto, A. J. Introducing ARTiMiS: A Low-Cost Flow Imaging Microscope for Microalgal Monitoring. Environ. Sci. Technol. 2024, 58 (30), 13540–13551. 10.1021/acs.est.4c01928.

(35) Patel, A.; Antonopoulou, I.; Enman, J.; Rova, U.; Christakopoulos, P.; Matsakas, L. Lipids Detection and Quantification in Oleaginous Microorganisms: An Overview of the Current State of the Art. BMC Chemical Engineering 2019, 1 (1), 13. 10.1186/s42480-019-0013-9.

(36) Liu, T.; Chen, Z.; Xiao, Y.; Yuan, M.; Zhou, C.; Liu, G.; Fang, J.; Yang, B. Biochemical and Morphological Changes Triggered by Nitrogen Stress in the Oleaginous Microalga Chlorella Vulgaris. Microorganisms 2022, 10 (3), 566. 10.3390/microorganisms10030566.

(37) Baroni, É.G.; Yap, K. Y.; Webley, P. A.; Scales, P. J.; Martin, G. J. O. The Effect of Nitrogen Depletion on the Cell Size, Shape, Density and Gravitational Settling of Nannochloropsis Salina, Chlorella Sp. (Marine) and Haematococcus Pluvialis. Algal Research 2019, 39, 101454. 10.1016/j.algal.2019.101454.

(38) Zaritsky, A.; Jamieson, A. R.; Welf, E. S.; Nevarez, A.; Cillay, J.; Eskiocak, U.; Cantarel, B. L.; Danuser, G. Interpretable Deep Learning Uncovers Cellular Properties in Label-Free Live Cell Images That Are Predictive of Highly Metastatic Melanoma. cels 2021, 12 (7), 733-747.e6. 10.1016/j.cels.2021.05.003.

