## Supporting Information for "Encoding Cell Phenotype from Label-Free Imaging Flow Cytometry with Unsupervised Deep Learning"

***Table S1:*** Feature threshold filter criteria to exclude “low-quality” events from VAE training and evaluation. Inclusion criteria select for “high-quality” events.

| **Feature** | **Inclusion Criteria** |
| --- | --- |
| Aspect Ratio | Value > 0.75 |
| Area | 4.5 < Value < 70 |
| Circularity (Hu) | Value > 0.85 |
| Centroid (X) | 27 < Value < 43 |
| Centroid (Y) | 27 < Value < 43 |
| Edge Gradient | Value > 0.28 |
| Edge Noise (Laplacian) | Value < 27 |
| Solidity | Value > 0.89 |
| Number of Objects in ROI | Value == 1 |

***Table S2:*** VAE model architecture detailed by layer.

| Layer | Element | Activation |
|---|---|---|
| Encoder: Layer 1 | Input (70x70x1 image) | |
| Encoder: Layer 2 | 2D Convolution (32 nodes, 5x5 kernel) | ReLU |
| Encoder: Layer 3 | 2D Convolution (16 nodes, 5x5 kernel) | ReLU |
| Encoder: Layer 4 | 2D Convolution (8 nodes, 3x3 kernel) | ReLU |
| Encoder: Layer 5 | Flatten | None |
| Encoder: Layer 6 | Dense (32 nodes) | ReLU |
| Latent Layers | Dense (2 latent dimensions) | None |
| Decoder: Layer 1 | Dense (28800 nodes) | ReLU |
| Decoder: Layer 2 | Reshape (60x60x8 image) | None |
| Decoder: Layer 3 | 2D Convolution Transpose (8 nodes, 3x3 kernel) | ReLU |
| Decoder: Layer 4 | 2D Convolution Transpose (16 nodes, 5x5 kernel) | ReLU |
| Decoder: Layer 5 | 2D Convolution Transpose (32 nodes, 5x5 kernel) | ReLU |
| Decoder: Layer 6 | 2D Convolution Transpose (1 node, 3x3 kernel) | Sigmoid |
| Decoder: Layer 7 | Output (70x70x1 image) | |

***Table S3:*** One-sided t-test evaluating whether one model (columns) demonstrated a lower RMSE value than a paired model (rows). For example: FCM 2D RF has an RMSE value significantly lower than all other models. Significance levels as follows: $p < 0.05$ (*), $p < 0.01$ (**), $p < 0.001$ (***). FT = Feature Table, VAE = Variational Autoencoder (LD1 and LD2), FCM = Flow Cytometry (SSC and FSC). LR = Linear Regression, RF = Random Forest Regression.

| | FT FD LR | FT FD RF | FT 1D LR | FT 2D LR | FT 2D RF | VAE 1D LR | VAE 2D LR | VAE 2D RF | FCM 1D LR | FCM 2D LR | FCM 2D RF |
|---|---|---|---|---|---|---|---|---|---|---|---|
| FT FD LR | | | | | *** | *** | *** | *** | | * | *** |
| FT FD RF | | | | | *** | *** | *** | *** | * | ** | *** |
| FT 1D LR | *** | *** | | | *** | *** | *** | *** | *** | *** | *** |
| FT 2D LR | *** | *** | | | *** | *** | *** | *** | *** | *** | *** |
| FT 2D RF | | | | | | | | | | | *** |
| VAE 1D LR | | | | | | | | | | | *** |
| VAE 2D LR | | | | | | | | | | | *** |
| VAE 2D RF | | | | | * | ** | * | | | | *** |
| FCM 1D LR | | | | | *** | *** | *** | * | | | *** |
| FCM 2D LR | | | | | *** | *** | *** | | | | *** |
| FCM 2D RF | | | | | | | | | | | |

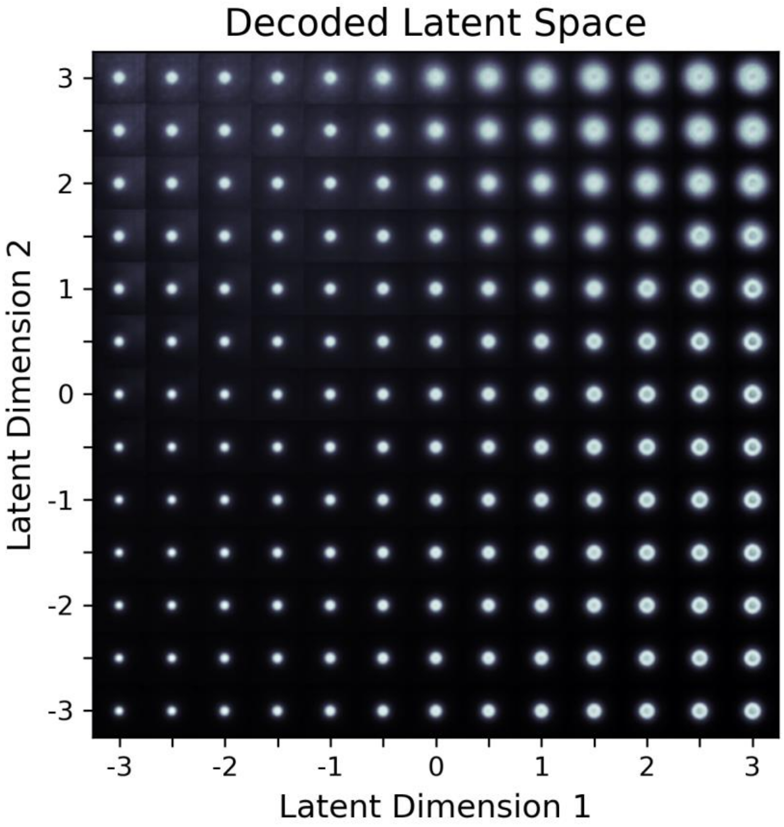


**Figure S1:** Latent space decoding from VAE model decoder output, visualizing archetypal images corresponding to specific coordinates in latent space.

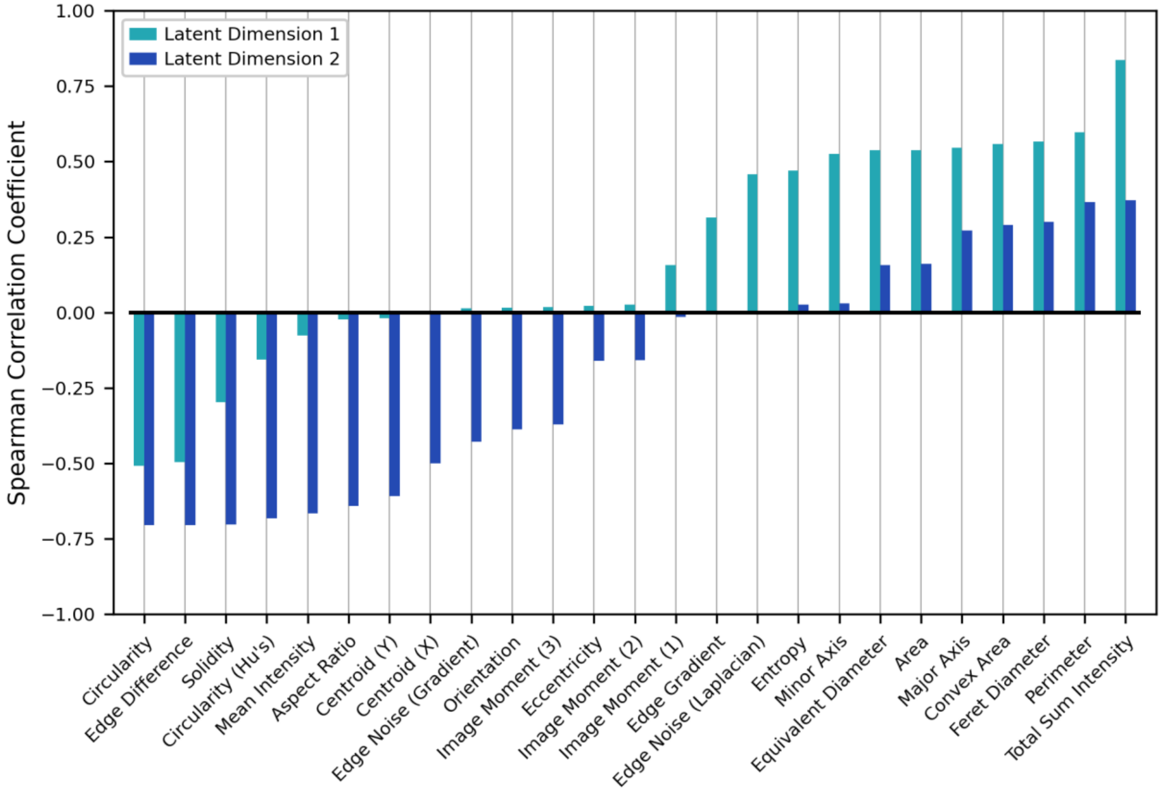


**Figure S2:** Spearman correlation coefficients indicating the relationships between individual microalgal morphological features and their respective VAE embedding latent dimension coordinates, evaluated on a single-cell basis. Greater magnitudes imply greater relatedness between a particular feature and a latent dimensional axis. For example, cell circularity appears to be encoded in both latent axes, and in the same direction (as latent coordinate increases, circularity decreases), whereas Aspect Ratio appears to be encoded only in LD2.
